# Fractal-Thermodynamic System Analogy and Complexity of Plant Leaves

**DOI:** 10.1101/2022.07.05.498782

**Authors:** M Vishnu, R Jaishanker

## Abstract

More precise measurements of the complexity of leaf shapes can open new pathways to understanding plant adaptation and resilience in the face of changing environments. We present a method to measure the complexity of plant leaf shapes by relating their fractal dimensions to topological entropy. Our method relies on ‘segmental fractal complexity’ and stems from a fractal-thermodynamic system analogy. The complexity of plant leaf shapes is an algebraic combination of the fractal dimensions of the components of the leaf-image system. We applied this method to leaf forms of 30 tropical plant species. While topological entropy is positively correlated with the leaf dissection index, it is an improvement over the leaf dissection index because of its ability to capture the spatial positioning of the leaf lamina, the leaf edges, and the leaf background. The topological entropy method is also an advancement over conventional geometric and fractal dimension-based measures of leaf complexity because it does not entail information loss due to the pre-processing and is perceptibly simple.

## 1. Introduction

The complexity of biological forms has fascinated mankind over the years. Early studies on complexity were primarily visual [1–4]. Quantitative morphometry emerged as an active research theme during the mid 20^th^ Century [5–7]. Morphometry has piggybacked on computer science to derive shape models [8,9]. From an information-theoretic point of view, the analysis of geometrical structures begins with Shannon’s work [10], expressing complexity as the amount of information (entropy) required to describe a phenomenon at a particular scale [11]. Entropy measures effectively describe complex patterns and have many applications in science and technology, ranging from medical imaging [12,13], remote sensing [14,15], security [16,17] and materials science [18,19].

Our current understanding of the morphological diversity of biological forms is strongly dependent on the effectiveness of available geometric, computational, and information methodologies in morphometrics. Approaches to clearly and completely describe the morphology of biological forms are still challenging [20,21]. Therefore, it is essential in design studies to find a more improved and automated method that minimizes information loss and time consumption. Plant leave shapes are highly diverse, and their morphology determines the capture and utilization of solar energy [22–24] and water [25,26]. Unravelling the physiological and evolutionary basis of the observed diversity of leaf shapes (complexity) is necessary for understanding the evolutionary history of plant physiology, reconstructing paleoclimatic conditions using fossil leaves [27,28], as well as the adaptive potential of contemporary plant species in the face of a rapidly changing climate and biosphere.

Few plant leaves possess shapes that can be adequately quantified by Euclidean geometry. Therefore, various indices of leaf shape have been developed, with varying levels of success. The ratio of the length of the primary and secondary lobe [29,30], leaf dissection index [31,32], leaf roundness index [33,34], degree of incision [35] are commonly used indices that depict leaf shapes. Despite the promising vista offered by digital imaging in leaf morphological studies [36–40], the investigations were disproportionately skewed to developing algorithmic and computational techniques for automated species identification than representing leaf shape. On the other hand, statistical analysis of leaf variables, landmark and outline (Elliptical Fourier/Eigenshape) based imaging techniques [41–45] of morphometry reduce information content. Neither leaf mensuration indices nor image-derived metrics accurately represent leaf shape [37,46]. The irregular shape of leaf lamina and edge-waviness add to the uncertainties and restrict the utility of conventional methods to represent leaf shapes accurately.

The edge waviness of plant leaves can be represented in a nonlinear sequence, wherein each position is defined by its predecessors. The iterative recursion that renders waviness to leaf edge qualifies them as natural fractals [47,48]. Digital technology facilitated the adoption of fractal measures to characterize structures in biology [49,50], geology [51], and medicine [52]. Fractal analysis of plant leaves for taxonomic studies [53,54] and the characterization and discrimination of plant leaves [55–58] are well researched. Notable among fractal-based plant leaf studies is Ampelography (identification and classification of grapevines) [59]. However, analysis of fractal dimensions of real-world objects (apparent fractality) has congenital shortcomings like issues with linear regression due to lack of sufficient data points, limited range, and effect of abundance and lacunarity (gaps and heterogeneity) [60]. Hence, there exists a need for a more inclusive complexity measure of plant leaf shapes. Here, we present a fractal-thermodynamic analogy that connects ecological and thermodynamic systems and use topological entropy to represent the complexity of plant leaf shape.

## 2. Thermodynamic analogy of fractal image systems

The thermodynamic understanding of fractal systems underpins the fractal behaviour of natural phenomena. The following section presents some general considerations on the thermodynamic analogy of a fractal image system. The analogy assumes importance when the energetics between a system and its environment are considered. It provides ground for the thermodynamic interpretation of a natural fractal system.

### 2.1. Fractal image systems

A Fractal image system is a part of the physical world defined in a 2-D plane upon which investigation is focused. Like the thermodynamic system, fractal systems are separated from the universal space by an external environment called surroundings through imaginary boundaries.

Based on the possible exchange of mass and energy, thermodynamic systems are classified as open, closed, and isolated [61]. For analysis, a fractal image system is constrained as a 2-D space, completely isolated from its environment. However, natural fractal systems are open thermodynamic and exchange matter and energy with their surroundings.

### 2.2. Characteristics of fractal image systems

Macroscopic characteristics such as mass, volume, energy, pressure, and temperature describe a thermodynamic system [62]. Fractal image systems also possess specific characteristics or properties to describe their state. The change of a thermodynamic system from an initial to a final state is known as a thermodynamic process. Since the natural fractal system is thermodynamic [63], state changes also occur in them with time. However, fractal image systems are static visual illustrations of the natural fractal world. They are the representatives of an instantaneous condition. Hence their intrinsic properties are invariant and they never undergo a process unless subsequent state changes are accounted for with new fractal images. Like thermodynamic systems, fractal image systems also have two broad properties: intensive properties and extensive properties.

#### 2.2.1. Extensive property

Any property which depends on the area or extent of the fractal image system is called extensive property. The properties like fractal measure (*K*) and entropy (*S*) are examples of extensive properties.

However, the value of the extensive property in the fractal image system is not always additive as in a thermodynamic system. Hence, the value of entropy (derived from the fractal dimension) for the composite system is not equal to the sum of the entropy of its components [60].

#### 2.2.2. Intensive property

Any property independent of the area or extent of the fractal image system is called intensive property. Intensive properties are scale-independent. Fractal dimension (*D*) and Area size (*SA*) [64] are examples of intensive properties.

#### 2.2.3. Normalized values

Analogous to the special type of intensive properties, specific value and molar value in the thermodynamic system [65], a few intensive normalized fractal properties exist in the fractal image system, viz. Normalized entropy 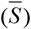, Normalized fractal dimension 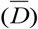 and Normalised Fractal measure 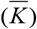 [66].

The ratio of the measured entropy (*S*) to the maximum entropy (*S*_*max*_) is called the Normalized entropy 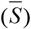. The maximum entropy is the logarithm of the total number of pixel elements in the fractal image system.

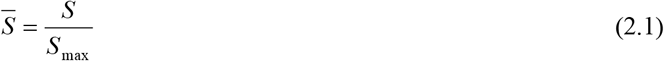

The ratio of the measured fractal dimension (*D*) to the maximum fractal dimension (*D*_*max*_) is called the Normalized fractal dimension 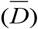. The maximum fractal dimension is the Euclidean dimension of a 2-D plane (i.e. 2).

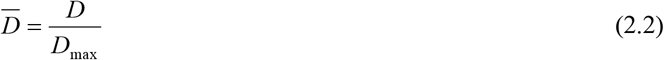

The scale-independent normalized entropy is equivalent to the normalized fractal dimension.

Normalized Fractal measure 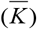 is the ratio of fractal measure (*K*) to the maximum fractal measure (*K*_*max*_) of the fractal image system. The maximum fractal measure has a value equal to the total area of the fractal image system in pixels.

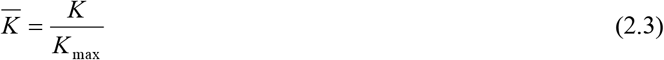

The analogy between the thermodynamic system and the fractal image system is summarised in table 1.

**Table 1.** A thermodynamic analogy of fractal image system.

| **Description** | **Thermodynamic system** | **Fractal image system** |
| --- | --- | --- |
| System | A definite quantity of matter or a region in space | A part of the physical world defined in a 2-D plane |
| Composition | The system, Boundary and Surroundings | The system, Boundary and Surroundings |
| Types | Open, Closed, Isolated | Isolated (static 2-D representation of a physical system) |
| Extensive Properties | Mass ($m$), Volume ($V$), Energy ($E$), Entropy ($S$) | Fractal measure ($K$), entropy ($S$) |
| Intensive properties | Pressure ($P$), temperature ($T$) | Fractal Dimension ($D$), Area size ($S_A$) |
| Derived properties | Specific volume ($v$), Specific internal energy ($\mu$), Molar volume ($\bar{v}$) | Normalised entropy ($\bar{S}$), Normalised fractal dimension ($\bar{D}$), Normalised fractal measure ($\bar{K}$) |

## 3. Materials and methods

Mature, healthy leaves of 30 flat-leaved plant species were collected from Trivandrum, Kerala, India. Morphological characterization of the leaves was carried out by capturing the full leaf images using a Digital scanner. The images were scaled to 1024 × 1024 pixels and converted into bitmap format (24-bit). The scaled colour images were transformed into grayscale, and Otsu’s threshold [67] was determined. The greyscale images were transformed into binary images using Otsu’s threshold. The fractal complexity of the entire leaf image system was analyzed by transforming the leaf image system into three discrete segments, namely ‘system’ (leaf lamina), ‘surroundings’ (the background), and ‘boundary’ (leaf edge) using the thermodynamic analogy. The ‘surrounding’ of the leaf image system is envisaged by removing the ‘leaf’ region from the leaf image system. Therefore, the leaf segments incorporate the waviness of the leaf edges. For analytical purposes, we consider the three segments as natural fractals. The representative images of the segments, ‘system’, ‘surrounding’ and ‘boundary’ are depicted in figure 1(a), figure 1(b), and figure 1(c), respectively.

**Figure 1.**
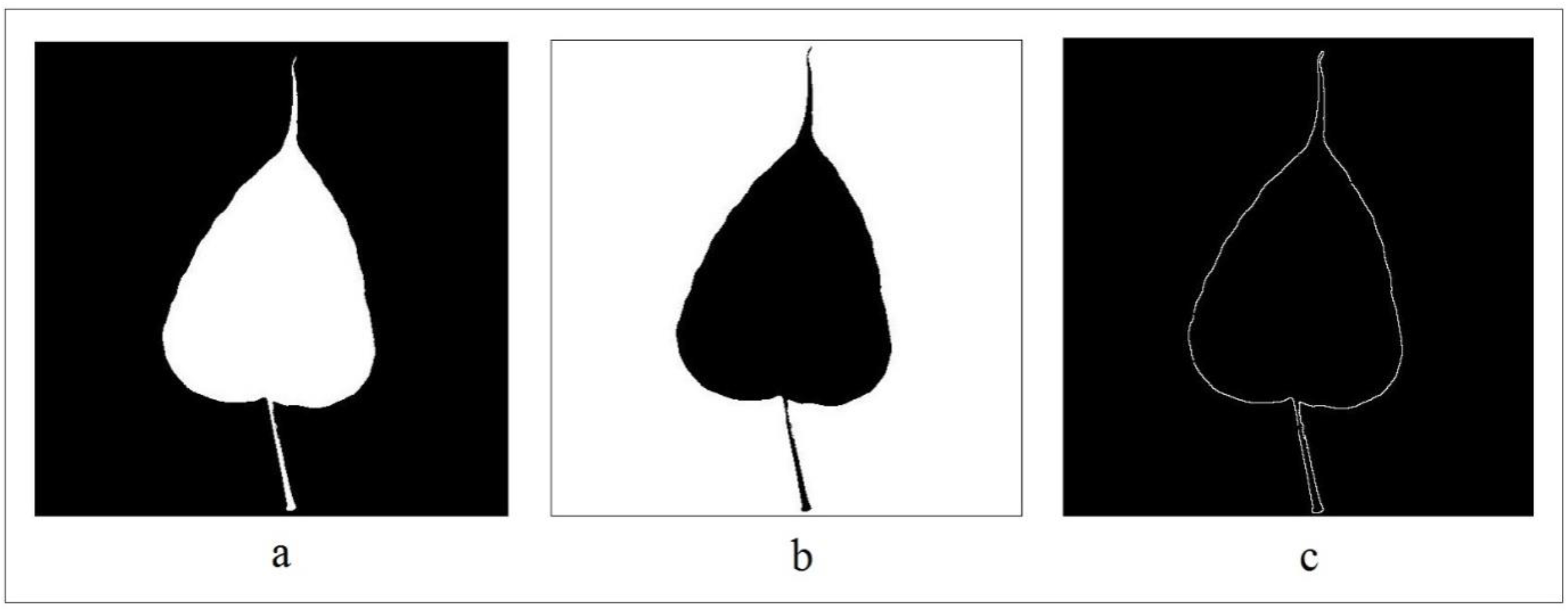
Representative image of (a) System, represents the image complement of original binary image (b) Surrounding, represents the original binary image and (c) Boundary, retrieved by extracting the perimeter pixels of the original binary image.

### 3.1. Fractal analysis

The fractal dimension of the threshold leaf image was evaluated using the Box Counting algorithm in Matlab® [68]. A regular mesh of boxes, composed of side length *δ* was superimposed on the threshold leaf image system. The number of boxes *N* needed to cover the characteristic fractal part (boxes that entirely cover the set and boxes which partially cover the set) was measured. The process was repeated for various box sizes *δ*, and the number of boxes *N(δ)* was computed.

A true/mathematical fractal set exhibits power-law behaviour of the form:

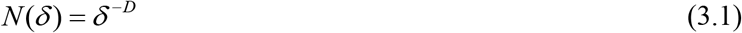

where *D* is the box-counting dimension.

However, a natural fractal set shows fractal properties only over a limited range of box sizes *δ*. Therefore, the local fractal dimensions were precisely determined from the logarithmic dependence of the number of boxes *N(δ)* and box size *δ*.

Local fractal dimension (*D*) was calculated as:

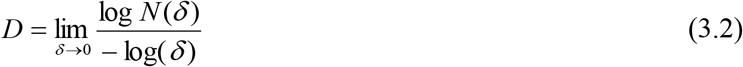

The local fractal dimensions computed were not constant for every box size. The actual fractal dimension of the fractal leaf image was calculated by taking the mean of at least six consecutive local fractal dimensions with the least standard deviation.

The three independent fractal dimensions *D*_*Leaf*_, *D*_*Background*_, and *D*_*Edge*_, were computed for the transformed segments, leaf lamina, the background, and the leaf edge.

### 3.2. Segmental fractal complexity

The fractal dimensions *D*_*Leaf*_, *D*_*Background*_, and *D*_*Edge*_ of the leaf images system describe the characteristics of the fractal leaf image parts, viz. leaf lamina, the background, and leaf edge, respectively.

A segmental fractal complexity (*D*_*ΣS*_) was calculated as a combination of three independent fractal dimensions.

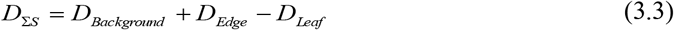

### 3.3. Entropy estimation

The Fractal dimension (*D*) and Renyi entropy (*R*) of a complex system were related to each other by the box-counting method [69,70].

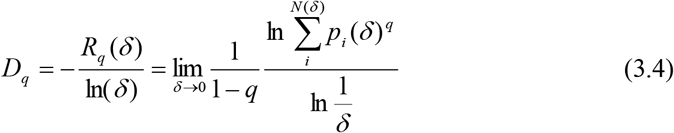

where *q* refers to the order of moment, *p*_*i*_*(δ)* is the probability of occurrence of fractal elements in the system, and *N(δ)* is the number of nonempty boxes of linear size *δ*.

The topological entropy or Hartley’s macro state entropy (*R*_*0*_) [64,71] of the system as a special case of equation (3.4) can be defined for the order *q = 0* as:

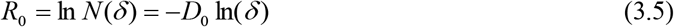

Two topological entropy, *S*_*L*_ and *S*_*ΣS*_ of the leaf images, were estimated from the traditional fractal dimension *D*_*Leaf*_, and segmental fractal complexity *D*_*ΣS*_, respectively, using equation (3.5).

A scale-independent normalized segmental fractal complexity 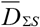 (normalized fractal dimension, 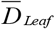 in traditional fractal analysis), which is equivalent to normalized topological entropy 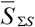 (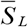 in traditional fractal analysis) was calculated using equations (2.1) and (2.2).

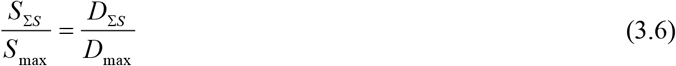

where *S*_*max*_ and *D*_*max*_ denote the maximum entropy and maximum segmental fractal complexity of the leaf image system, respectively.

### 3.4. Leaf dissection index

Leaf dissection index (*LDI*) is a dimensionless number that captures the degree of complexity of leaf shapes. *LDI* of the leaf image system was computed as the ratio of the leaf’s outline perimeter to the square root of the leaf area [32,72].

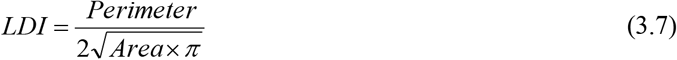

The *S*_*L*_ and *S*_*ΣS*_ of the leaf images were correlated with the estimated *LDI*.

## 4. Results

The leaves of 30 plant species collected from Trivandrum, Kerala, India, show remarkable morphological diversity (figure 2). A description of the shape of each leaf is presented in table 2.

**Table 2.** Taxonomic and morphological description of plant leaves collected for the study.

|  | **Species** | **Genus** | **Family** | **Leaf Type** |
| --- | --- | --- | --- | --- |
| 1 | *Bauhinia purpurea* | Bauhinia | Fabaceae | Simple and deeply lobed |
| 2 | *Artocarpus hirsutus* | Artocarpus | Moraceae | Simple |
| 3 | *Clidemia hirta* | Clidemia | Melastomataceae | Simple and toothed |
| 4 | *Piper nigrum* | Piper | Piperaceae | Simple |
| 5 | *Lagerstroemia speciosa* | Lagerstroemia | Lythraceae | Simple |
| 6 | *Annona muricata* | Annona | Annonaceae | Simple |
| 7 | *Terminalia bellirica* | Terminalia | Combretaceae | Simple |
| 8 | *Macaranga peltata* | Macaranga | Euphorbiaceae | Simple |
| 9 | *Merremia vitifolia* | Merremia | Convolvulaceae | Simple, deeply lobed and toothed |
| 10 | *Psidium guajava* | Psidium | Myrtaceae | Simple |
| 11 | *Artocarpus heterophyllus* | Artocarpus | Moraceae | Simple |
| 12 | *Terminalia arjuna* | Terminalia | Combretaceae | Simple |
| 13 | *Xanthophyllum flavescens* | Xanthophyllum | Polygalaceae | Simple |
| 14 | *Ficus exasperate* | Ficus | Moraceae | Simple |
| 15 | *Acacia mangium* | Acacia | Fabaceae | Simple |
| 16 | *Syzygium jambos* | Syzygium | Myrtaceae | Simple |
| 17 | *Hydnocarpus pentandrus* | Hydnocarpus | Achariaceae | Simple |
| 18 | *Tecoma stans* | Tecoma | Bignoniaceae | Simple and toothed |
| 19 | *Mangifera indica* | Mangifera | Anacardiaceae | Simple |
| 20 | *Monoon longifolium* | Monoon | Annonaceae | Simple |
| 21 | *Manilkara zapota* | Manilkara | Sapotaceae | Simple |
| 22 | *Ficus religiosa* | Ficus | Moraceae | Simple |
| 23 | *Acacia auriculiformis* | Acacia | Fabaceae | Simple |
| 24 | *Carica papaya* | Carica | Caricaceae | Simple and deeply lobed |
| 25 | *Hibiscus cannabinus* | Hibiscus | Malvaceae | Simple, deeply lobed and toothed |
| 26 | *Albizia odoratissima* | Albizia | Fabaceae | Bipinnate |
| 27 | *Murraya koenigii* | Murraya | Rutaceae | Pinnate |
| 28 | *Moringa oleifera* | Moringa | Moringaceae | Tripinnate |
| 29 | *Azadirachta indica* | Azadirachta | Meliaceae | Pinnate and toothed |
| 30 | *Phyllanthus emblica* | Phyllanthus | Phyllanthaceae | Pinnate |

**Figure 2.**
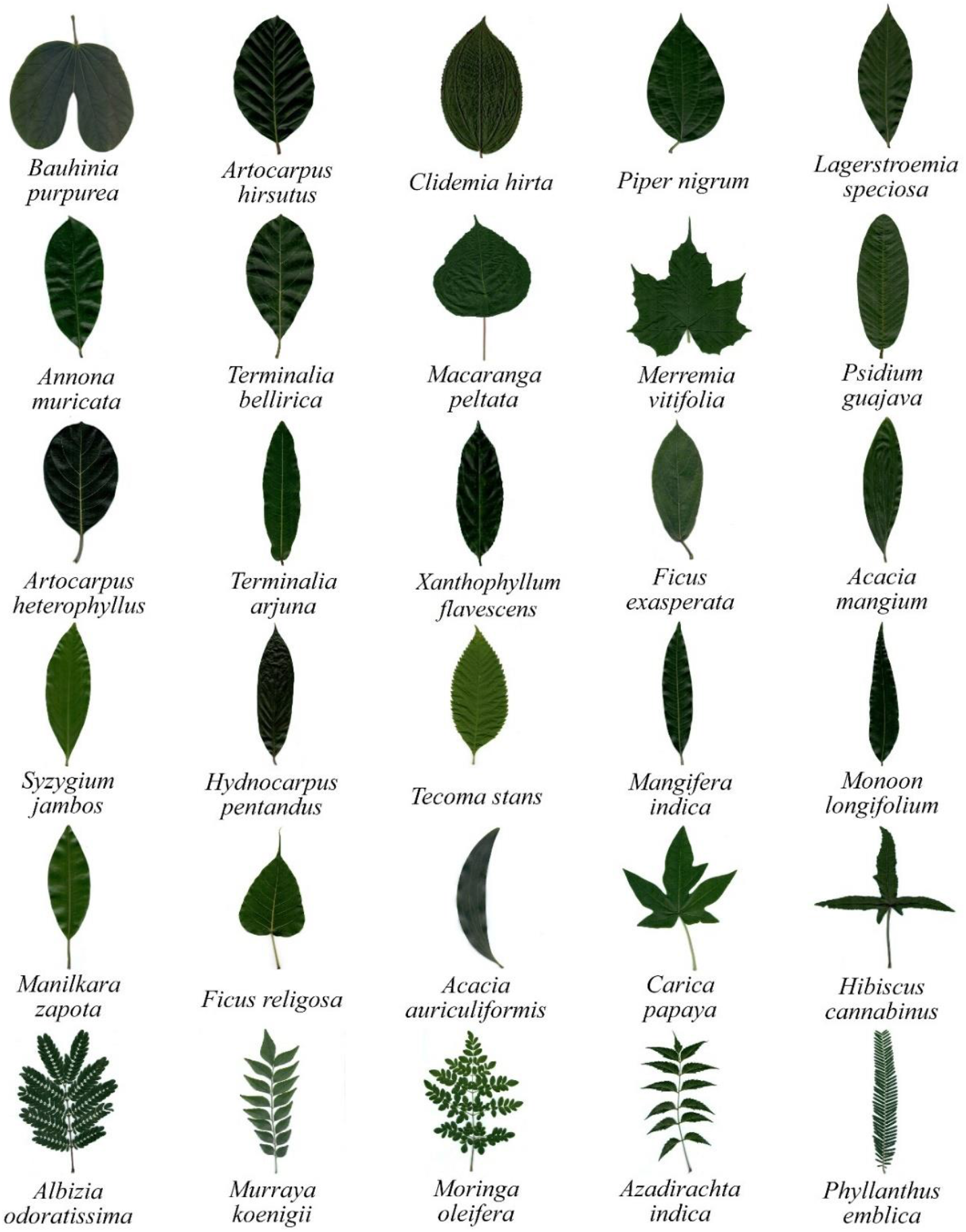
Morphological diversity of the plant leaves studied.

### 4.1. Traditional fractal analysis of leaves

The traditional fractal analysis of the leaf images using the box-counting technique is presented in table 3. The structural complexity of the leaves is described by the fractal dimension of the leaf (*D*_*Leaf*_), topological entropy (*S*_*L*_), and normalized topological entropy (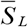). *D*_*Leaf*_ in the study varies between 1.568 and 1.946. *S*_*L*_ varies between 10.869 and 13.490. The leaf of *Azadirachta indica* showed the lowest *S*_*L*_ (10.869) and that of *Artocarpus hirsutus*, the highest (13.490).

**Table 3.**
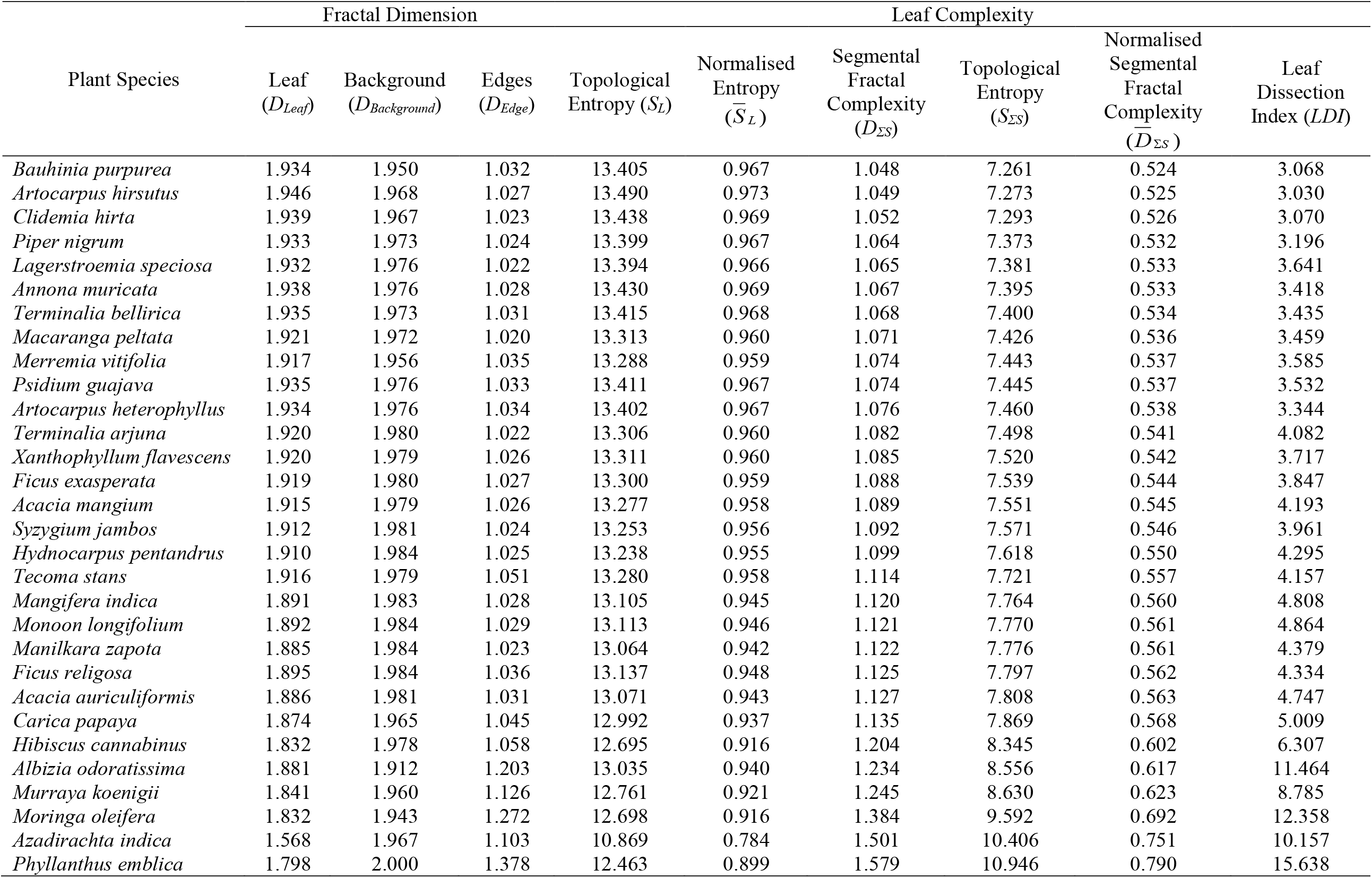
Fractal dimensions and Leaf Complexity of 30 plant species studied.

*S*_*L*_ does not show significant variation to discriminate the leaf forms. Most of the leaves have comparable *S*_*L*_. The leaves of *Albizia odoratissima, Murraya koenigii, Moringa oleifera, Azadirachta indica*, and *Phyllanthus emblica* are pinnately compound. However, their *S*_*L*_ values were lower than that of simple leaves. The scale-independent 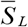 also showed less variation among the studied leaf samples (*μ* = 0.946, *σ* = 0.036) (figure 3). A maximum *LDI* of 15.638 and a minimum of 3.030 were obtained for the leaves of *Phyllanthus emblica* and *Artocarpus hirsutus*, respectively (table 3). The inverse relation between *S*_*L*_ and *LDI* is depicted in figure 4.

**Figure 3.**
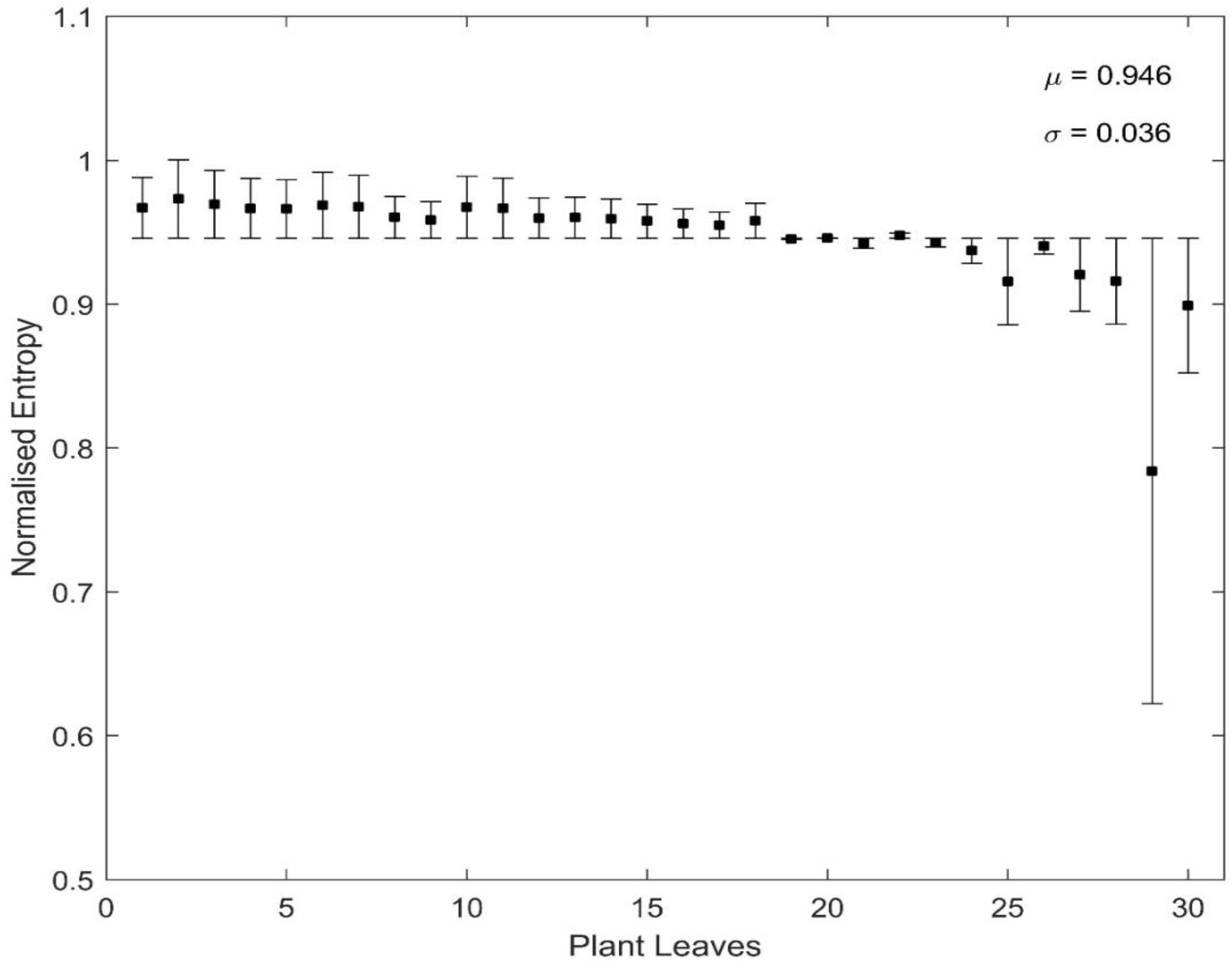
Variations of the normalised topological entropy from traditional fractal analysis among 30 studied plant leaves. The species numbering corresponds with that in table 2.

**Figure 4.**
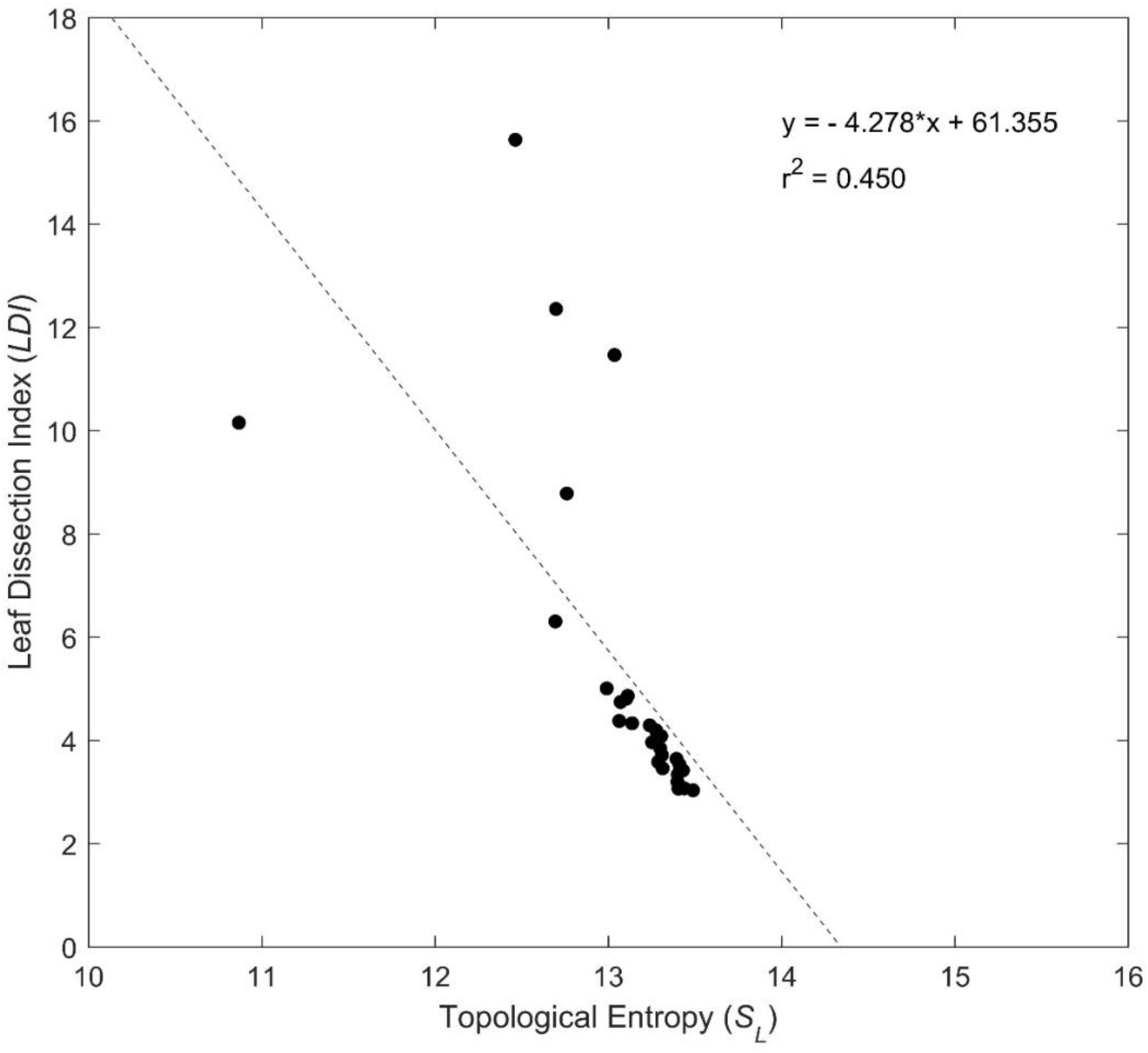
Correlation between Topological entropy, calculated by the traditional box-count analysis and Leaf dissection index for leaves of 30 plant species.

### 4.2. Segmental fractal complexity of leaf image system

The segmental fractal complexity *(D*_*ΣS*_) of the leaf image systems are presented in table 3. *D*_*ΣS*_ varies between 1.048 and 1.579. The typical topological entropy (*S*_*ΣS*_) of the leaf image system varies between values 7.261 and 10.946. The highest and lowest *D*_*ΣS*_ were recorded for *Phyllanthus emblica* (1.579) and *Bauhinia purpurea* (1.048), respectively (table 3).

*D*_*Leaf*_ value was highest (1.946) for *Artocarpus hirsutus* and lowest (1.568) for *Azadirachta indica* (table 3). The highest and lowest values of *D*_*Edge*_ were observed for the leaves of *Phyllanthus emblica* (1.378) and *Macaranga peltata* (1.020), respectively. However, *D*_*Background*_ values were comparable for all leaf image systems. The majority of the studied leaves were simple, and their *S*_*ΣS*_ ranged between 7.261 and 7.808. Deeply lobed, simple leaves of *Carica papaya* and *Hibiscus cannabinus* recorded *S*_*ΣS*_, 7.869 and 8.345, respectively. The *S*_*ΣS*_ of the pinnately compound leaves of *Albizia odoratissima, Murraya koenigii, Moringa oleifera, Azadirachta indica*, and *Phyllanthus emblica* were the highest and ranged between 8.556 and 10.946 (table 3).

The scale-independent normalized segmental fractal complexity (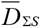) was higher for the compound leaf of *Phyllanthus emblica* (0.790) and lower for the simple leaf of *Bauhinia purpurea* (0.524). Variations in the 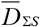 follow the same trend as *D*_*ΣS*_. Figure 5 illustrates very high, positive correlation (*r =* 0.94) between the *S*_*ΣS*_ and *LDI*.

**Figure 5.**
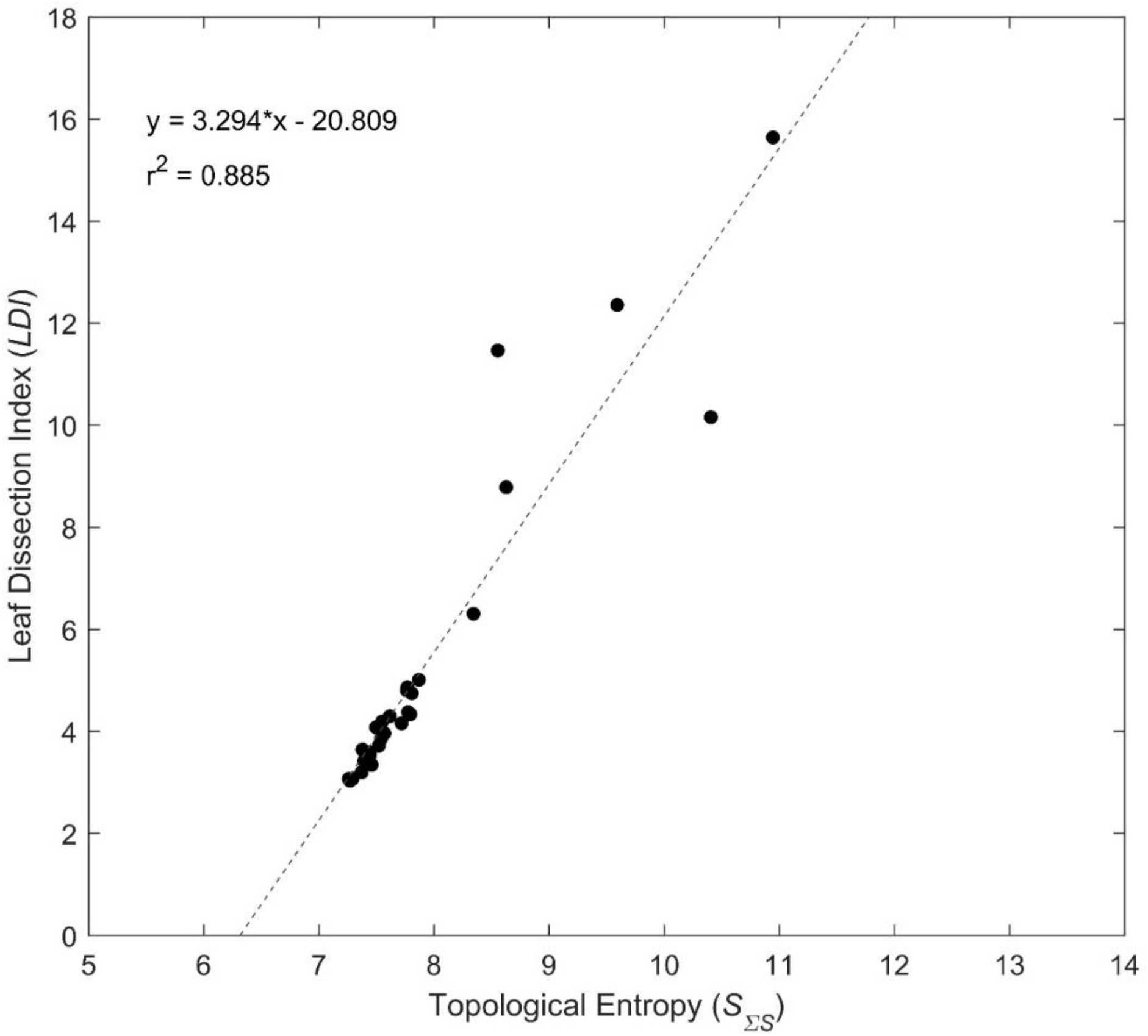
Correlation between Topological entropy and Leaf dissection index for leaves of 30 plant species.

### 4.3. Theoretical extremes of plant leaf complexity

The *D*_*ΣS*_ of the leaf image system was calculated by computing the fractal dimensions, *D*_*Leaf*_, *D*_*Background*_ and *D*_*Edge*_. The method was applied to selected natural leaf image systems. However, the variations in the complexity of the natural leaf shapes hardly advance into the theoretical extremes of *D*_*ΣS*_.

Based on the space-filling capacity of the fractal elements in the leaf image system, we define three hypothetical angiosperm leaf shapes to illustrate the theoretical extremes of *D*_*ΣS*_ of the leaf image system. Figure 6(a) describes the leaf shape having maximum leaf space-filling capacity. The maximum leaf coverage can be attained only for square shape leaf as the space of interest is a box. The imaginary edge or boundary part is also a square curve. Similarly, the least leaf coverage in figure 6(b) is for a fractal image system with maximum ‘background’ coverage. An exceptional leaf shape having a maximum space-filling capacity for all three components (leaf lamina, the background, and leaf edge) of the leaf image system is described in figure 6(c). This kind of space-filling fractal shape is common in fractal geometry. Peano curve is one such fractal shape having maximum space-filling capacity [73]. The imaginary shape in figure 6(c) was deduced from knowledge of the Peano curve in a square region. The extreme values of the *D*_*ΣS*_ and *S*_*ΣS*_ of the theoretical leaf image systems were calculated using the box-counting method. The results are presented in table 4. *D*_*Leaf*_ of the hypothetical leaf shapes in table 4 represents the leaf complexity using the traditional fractal dimension. The highest *D*_*ΣS*_ is recorded for figure 6(b) and figure 6(c). Figure 6(a) showed the lowest *D*_*ΣS*_. The *D*_*ΣS*_ of hypothetical leaf shapes increases with complexity.

**Table 4.** Complexity of the three extreme leaf shapes using box-count analysis.

| Fractal Image System | Leaf<br>($D_{Leaf}$) | Background<br>($D_{Background}$) | Edges<br>($D_{Edge}$) | Segmental Fractal Complexity<br>($D_{\Sigma S}$) | Topological Entropy<br>($S_{\Sigma S}$) |
| --- | --- | --- | --- | --- | --- |
| Figure 6(a) | 2 | 1 | 1 | 0 | 0 |
| Figure 6(b) | 1 | 2 | 1 | 2 | 13.863 |
| Figure 6(c) | 2 | 2 | 2 | 2 | 13.863 |

**Figure 6.**
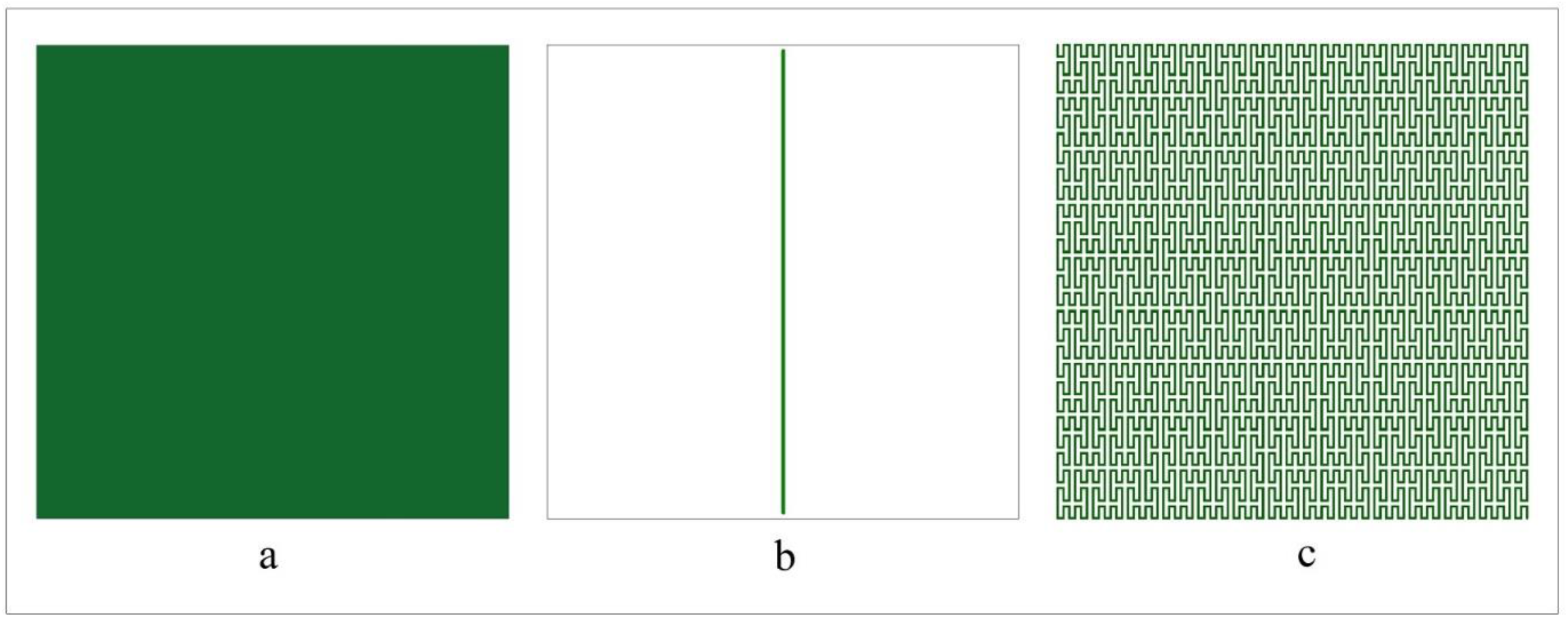
Three extreme hypothetical leaf shapes: (a) Maximum leaf space coverage (b) Minimum leaf space coverage (c) Maximum leaf-background-edge coverage.

## 5. Discussion

Leaf morphology is an inheritable plant trait. It influences light absorption, sap transport, and photosynthesis [74]. Plant leaves are classified as simple and compound. Simple leaves contain only a single leaflet. Multiple leaflets organized along the rachis form compound leaves [75].

Combining simple and compound leaves with entire, serrated and lobed marginal features offers enormous complexity. Plants optimize leaf morphology to increase energy exchange efficiency, maximize carbon assimilation [76,77], apportion resources for growth [78], reproduction [79], and resistance [80]. Since phytohormones, genetic and environmental factors govern leaf developmental traits [81,82], biochemical and molecular pathways have been predominantly used to understand them. Here we leverage fractal geometry to explain leaf shape.

### 5.1. Fractal optimization of leaf forms

Primitive terrestrial flowering plants had broad, simple leaves and evolved to compound leaves over time [83,84]. This evolutionary adaptation enables terrestrial plants to absorb light uniformly and optimize photosynthesis [85,86]. However, leaf size is associated with the architectural structure of the plant. The size of leaves depends on plant height [87] and its life span [88,89]. Plants have evolved to arrive at an optimal balance between leaf size and the number of leaves [90]. Limiting the size and number of leaves affects the growth of plants. From a developmental perspective, the evolution of plant leaves from simple to compound follows a fractal pattern, thereby reducing leaf overlap and enabling them to receive adequate sunlight without compromising the surface area.

### 5.2. Fractal analysis

At a fine scale, the area covered by the plant leaves is strictly two dimensional. However, their edges behave like fractals. Moreover, leaf area and edge assume importance if the flow of materials between the leaf and the environment are considered [91,92]. We consider the plant leaf a natural fractal [93,94]. Since the geometric properties of natural fractals are restricted over a limited scale [95], fractal dimension approximation of natural leaves seems impractical over the complete range of scales. However, the variation of fractal dimension over the limited range of scales indicates the changes in the information as landscape changes during succession [96], and aggregate structure changes in industrial applications [97]. The challenges of the limited range of fractal property of natural fractals and the lack of sufficient data points for linear regression [60,98] were overcome by using the mean of at least six consecutive local fractal dimensions with the least standard deviation. Completely covered grids in a 2-D space result in a box-shaped leaf with a fractal dimension of 2, and the least filled grids result in a linear-shaped leaf with a minimum fractal dimension. The information entropy quantified from the fractal dimensions of the threshold leaf patterns indicates only the presence or absence of the fractal leaf constituents. It does not convey anything about the pixel characteristics. Further, this topological entropy [64] depends only on the bulk and boundary of the leaf shape and is sensitive to the linear scale of leaf image systems. The normalized topological entropy of the leaf images is a scale-independent entropy measure that is equivalent to the corresponding normalized fractal dimension.

The traditional box-counting algorithm derived fractal dimensions (table 3) rely only on the space-filling capacity of constituent elements and is area dependent. A study of *D*_*Leaf*_ presented in table 3, and the leaf images in figure 2 reveals the latent incongruence. The *S*_*L*_ of deeply lobed, pinnately compound leaves of *Albizia odoratissima, Murraya koenigii, Moringa oleifera, Azadirachta indica*, and *Phyllanthus emblica*, were lower than that of the simple leaves of *Piper nigrum, Lagerstroemia speciosa, Annona muricata, Terminalia bellirica* and *Macaranga peltata. S*_*L*_ derived from the *D*_*Leaf*_ involves only on the bulk of the leaf and is independent of its perimeter. Hence, it is a direct consequence that the *S*_*L*_ is negatively correlated with *LDI*. Although the conventional fractal geometric approach effectively describes natural shapes [99,100], they have limited by their inability to discriminate shapes [101,102]. Box counting studies integrating pattern search algorithms to address these shortcomings of the traditional box-counting method are reported [103]. However, they remain confined to the bulk of the leaf [103].

The fractal dimension of the composite objects takes the largest dimension of its subsets [104]. Previous studies have focused on the fractal analysis of leaves either along their outline or through their bulk. The *D*_*ΣS*_ (table 3) introduced here is an improved complexity measure that comprises discrete fractal dimensions *D*_*Leaf*_, *D*_*Background*_ and *D*_*Edge*_. The highest *D*_*ΣS*_ was observed in the pinnately compound leaf of *Phyllanthus emblica* and the lowest in the simple leaf of *Bauhinia purpurea*. The apparent range of *S*_*ΣS*_ is low in the studied leaf samples. A few of them are comparable. However, tiny deviations in the box-counting fractal analysis are informative and account for substantial changes [105,106]. The extremal complexity patterns of hypothetical angiosperm leaves described in figure 6(a), figure 6(b), and figure 6(c) seldom occur in the real world. The extreme narrow leaf (figure 6(b)) and the Peano curve-shaped leaf (figure 6(c)) represents the maximum complexity by *D*_*ΣS*_. The *D*_*ΣS*_ of the studied leaves reveals that the complexity is lower for simple leaves and increases with lobed leaves and pinnately compound leaves.

The strong positive correlation between *S*_*ΣS*_ and *LDI* reflects the direct causal links between leaf complexity, dissection and serration features of plant leaves. Although an important measure of the complexity of leaves, *LDI* can sometimes be misleading. The *LDI* of an un-dissected narrow leaf may be the same as that of a dissected leaf with moderately effective width, higher than that of an un-dissected broader leaf, or lower than that of a relatively wider small dissected leaf. Further, *LDI* does not relate independently to the bulk and boundary of the leaves and fails to capture the spatial positioning of leaf lobes and leaflet arrangements. *D*_*ΣS*_ addresses these limitations of *LDI*. It captures the dissectional property and the spatial positioning of leaf lobes and leaflet arrangements.

*D*_*ΣS*_ outperforms traditional and geometric morphometrics in more than one way. Traditional morphometrics relies on linear distance between predetermined points and statistical approaches to analyze shapes. It results in the loss of geometric information while converting coordinate points into linear distances [107]. Moreover, the linear metric correlates with the size, which leads to redundancy in the measurements [108]. Unlike traditional morphometrics, *D*_*ΣS*_ is simple and does not involve any statistical analysis.

Geometric morphometrics is tedious and time-consuming. In the landmark approach, each shape studied must have unique landmarks. The inclusion of more landmarks will be redundant and affect the statistical analyses. In contrast to the landmark approach, the outline-based approach uses semi-landmarks. The outline is quantified using Elliptical Fourier and Eigenshape analysis. This method performs suboptimally with pointed shapes [108]. *D*_*ΣS*_ offers a unique numeric metric of shape. Devoid of statistical techniques, *D*_*ΣS*_ is free from any information loss. It captures the bulk and spatial positioning of every elemental shape descriptor in 2-D.

It is challenging to fully describe the complexity of Leaf forms [109]. The goal of science is to unravel the challenges put forward by the design patterns of complex systems like plant leaves. Although complex shapes stem from basic rules [110,111], deducing behaviour from such rules is difficult. The complexity bestowed by the self-organizing capability of individual *agents* is integral to the survival and progression of the system. It imparts flexibility and improves adaptive resilience capability. Understanding complexity using entropy, geometric, and computational approaches opens the pathway to describing complex natural systems’ formation, prediction, approximation, simulation, and environmental adaptation.

## 6. Conclusions

We quantify the complexity of plant leaf shapes based on an intuitive thermodynamic formalism of fractal image systems. The complexity of plant leaf shapes is objectively represented by relating fractal dimensions of leaf images to topological entropy. The segmental fractal complexity introduced in this paper overcomes information loss issues of traditional and geometric morphometric techniques. Understanding complexity in plant leaf shapes opens pathways to elucidate plant evolution and adaptive resilience. One day, it may help genetically engineer optimal leaf shapes to increase photosynthetic efficiency, crop yields, and carbon sequestration.

## Acknowledgement

We thank Dr. Saji Gopinath, Vice-Chancellor, Kerala University of Digital Sciences, Innovation and Technology, for providing all necessary support to carry out the study. We further gratefully acknowledge the support of Dr. Samraat Pawar, Imperial College, UK, and Dr. Sasidevan Vijayakumar, Cochin University of Science and Technology, India for detailed comments that led to a substantial improvement of the manuscript.

